# Resting-state fMRI correlations: from link-wise unreliability to whole brain stability

**DOI:** 10.1101/081976

**Authors:** Mario Pannunzi, Rikkert Hindriks, Ruggero G. Bettinardi, Elisabeth Wenger, Nina Lisofsky, Johan Martensson, Oisin Butler, Elisa Filevich, Maxi Becker, Martyna Lochstet, Ulman Lindenberger, Simone Kühn, Gustavo Deco

**Author notes:** Corresponding author, mail.

## Abstract

The functional architecture of spontaneous BOLD fluctuations has been characterized in detail by numerous studies, demonstrating its potential relevance as a biomarker. However, the systematic investigation of its consistency is still in its infancy. Here, we analyze both the within- and between-subject variability as well as the test-retest reliability of resting-state functional connectivity (FC) estimates in a unique data set comprising multiple fMRI scans (42) from 5 subjects, and 50 single scans from 50 subjects. To this aim we adopted a statistical framework enabling us to disentangle the contribution of different sources of variability and their dependence on scan duration, and showed that the low reliability of single links can be largely improved using multiple scans per subject. Moreover, we show that practically all observed inter-region variability (at the link-level) is not significant and due to the statistical uncertainty of the estimator itself rather than to genuine variability among areas. Finally, we use the proposed statistical framework to demonstrate that, despite the poor consistency of single links, the information carried by the whole-brain spontaneous correlation structure is indeed robust, and can in fact be used as a functional fingerprint.

## 1 Introduction

Neuroimaging techniques allow us to deal non-invasively with two main principles of brain functioning: segregation and integration. Relationships between segregate regions can be described at different scales, with different techniques, and the strengths of these relationships can help us to understand their integrative roles. Some methods help to describe the physical wiring between the brain regions (e.g., diffusion tensor imaging, tractography, etc), while others quantify the functional relationship between regions’ activity (Friston [2011]). To date, one of the most widely adopted techniques used to characterize the functional structure (also referred to as functional connectome) has been resting-state functional magnetic resonance imaging (rs-fMRI). Resting-state is commonly defined as the condition in which the participant is not performing any overt task, but lies still in the scanner (with eyes closed or fixating on a cross on a screen) while not focusing on any particular thought or sensation (see e.g., Biswal et al. [1995] or more recent Zuo and Xing [2014]). The method is based on the quantification of local changes in blood oxygenation through the use of the so-called blood-oxygen level-dependent (BOLD) signal (Ogawa et al. [1990]), that have been demonstrated to partially reflect underlying neural activations (Logothetis et al. 2001, Logothetis 2008, Magri et al. 2012). Functional connectivity (FC) between different regions of interests (ROIs) is then quantified with measures of statistical dependencies between such changes in different brain regions, with the Pearson correlation coefficient being the most commonly used (see e.g. Friston [2011]).

Resting-state functional connectivity (rs-FC) has already been adopted to differentiate between subjects Finn et al. [2015] and groups, either coming from healthy or pathological populations (see for example Rosazza and Minati [2011] for a review and references therein), or between different brain states (see for example the case of learning in Guerra-Carrillo et al. [2014] and references in there). The advantages of this method lie on its spatial resolution, speed and completeness (Logothetis [2008]). In fact, by using a resting-state fMRI scan of about 5 minutes, it is possible to obtain a large-scale description of the functional relationships between all brain areas. Those advantages make this technique potentially very powerful, even considering that it measures neural activity only indirectly through the BOLD signal (Logothetis [2008]). The unrestricted nature of the resting-state experiments could in fact mirror a wide range of cognitive states and operations (Christoff et al., 2009; Richiardi et al., 2011; Hurlburt et al., 2015).

Interestingly, functional connectome studies show a differential pattern of findings: on the one hand they show a very stable architecture of correlated spontaneous activity, on the other hand they indicate a high variability in the functional structure, with temporal dynamics ranging from less than one second (Mitra et al. [2015])), to days (Anderson et al. [2011]; Laumann et al. [2015]. According to the current literature, a crucial factor influencing the stability of the resting-state FC is scan duration. The most common acquisition time is 5–10 min, even though recent evidence indicates the importance of using much longer scans to obtain reliable FC estimates (Anderson et al. [2011]; Birn et al. [2013]; Hacker et al. [2013]; Laumann et al. [2015]). A question that has both theoretical and practical relevance is how much data we need to accurately and reliable estimate the FC of a single subject (Birn et al. [2013]; Laumann et al. [2015]; Finn et al. [2015]). One of the main objectives of our study is to reconcile these apparently conflicting aspects of resting-state FC.

The development of biomarkers derived from resting-state BOLD-fMRI scans that are able to characterize the functional architecture of individual brains is important for cognitive as well as for clinical neuroscience. For a biomarker to be successful, it has to be reliable; as such, two conditions must be met: on one hand, it should be stable for the same subject (or condition) across different sessions, whereas on the other hand it should substantially vary over different subjects (or conditions). The second requirement ensures that the biomarker is selective for the variable of interest, and could thus be used to effectively discern between different subjects or conditions. The principle behind the two above mentioned criteria suggests a rather straightforward way to quantify the reliability of a potential biomarker, namely by comparing the within-subject (-condition) variability with the between-subject (condition) variability. If these two types of variability can be described through the use of a normal variable, an index commonly adopted to measure this ratio is the intra-class correlation coefficient (ICC) a measure widely used in the psychological sciences to assess test-retest reliability (Shehzad et al. [2009]; Zuo and Xing [2014]). We will use the ICC as our main tool in assessing the reliability of resting-state FC.

Although numerous studies have been devoted to characterize the functional architecture of spontaneous BOLD-fMRI fluctuations, the test-retest reliability of functional indices has begun to be addressed only recently (Anderson et al. [2011]; Birn et al. [2013]; Hacker et al. [2013]; Zuo and Xing [2014]). From the results reported in the literature, one of the main findings is that test-retest reliability of functional indices between regions of interest (ROIs), as quantified by the intra-class correlation (ICC), seems to strongly vary over brain regions and over pairs of brain regions (for link-based indices). What has not been made explicit in previous studies, however, is an analysis of the variability of the reliability measures themselves. Indeed, reported variation of reliability has been interpreted to reflect differences in the reliability of the functional indices, without taking into consideration the statistical uncertainty due to finite sample in the estimates of the ICC.

Within the context of resting-state BOLD-fMRI, where the number of subjects and the number of scans by subject are usually limited, the variance of ICC estimators can be very high. Assessment of the variance of the estimated ICC is particularly relevant for investigating its heterogeneity over regions, links, and networks as done in Zuo and Xing [2014]. In fact, a proper assessment of the ICC variability was lacking in the above-cited studies, and as such its claimed that heterogeneity has still to be demonstrated.

In the present study, we replicated most of the analyses presented in Shehzad et al. [2009]; Birn et al. [2013]; Zuo and Xing [2014]; Laumann et al. [2015], as indeed they are pioneers in the analysis of both FC variability and reliability. In particular, we investigate the test-retest reliability of resting-state FC-fMRI, and its variability. To this aim, we use fMRI to measure the resting-state activity in a group of 6 participants, each of them scanned 50 times, which allowed assessing the intersession (session-to-session) reliability.

The paper is divided into three main sections, each one aimed at answering different questions: *In the first section*, we briefly present the data to provide the reader with an intuition of the variability and reliability of FC. *In the second section*, we analyze the variability and reliability of functional connectivity at the link level; the former estimated by the standard deviation of the correlation coefficients between ROIs, the latter through the ICC. We analyze how a finite number of samples influences both the variability and the reliability of the functional connectivity estimates. We conclude the first section showing that it is not possible to statistically differentiate between links based on their ICC value, as all the correlation coefficients can be described, as a first approximation, with a unique, low ICC value (≈0.2). As it has been suggested before (see e.g. a nice schematic resume Fornito et al. [2010]) that different parcellations can influence the resulting FC estimates, we repeat our analysis using two different parcellations: one based on anatomy, (AAL, Tzourio-Mazoyer et al. [2002]), and one based on functionality (Shen et al. [2013]). We focus on characterizing and quantifying the nature of variability observed in empirical functional connectivity by decomposing it into the variability due to finite-sample statistical fluctuations and into variability that is likely due to real dynamic changes in the strength of the functional connections. We systematically analyze the behavior of these variability factors both for different scan durations and for multiple sessions. *In the last section* we analyze the reliability of the whole FC matrix. For this purpose, we compare the FC matrices obtained in different sessions both within- and between-subject, showing that the complex information contained in those matrices is much more stable than the correlations between individual pairs of ROIs. Indeed, by means of a general linear model, we solve the apparent contradiction of low link-wise reliability and stability of the whole-brain FC matrix.

## 2 Materials and Methods

### 2.1 Data acquisition and pre-processing

Fifty eight participants were recruited. Eight of the participants volunteered to be included in the longitudinal part of the study in which they were scanned 40–50 times over the course of 6 months (2 male, mean age 29, SD= 2.6, range: 24–32). Two of the participants (one male, one female) did not find the time to continue with the study and had to be excluded from further analysis. We had to exclude even the last male participant, who, in contrast to the instruction received, tried to apply relaxation exercise during the scan which largely influenced the measure (see figure S3 in the supplementary material). The other fifty participants (all female, mean age 24, SD=3.1, range: 18–32) were part of another study that was conducted during the same period of time and underwent scanning with the same MRI sequences only once. According to personal interviews (Mini-International Neuropsychiatric Interview, Margraf [1994]) the participants to the longitudinal study were free of psychiatric disorder and had never previously suffered from a mental disease. The other participants were asked on the phone during recruitment whether they ever had a psychiatric disease and negated that. Other medical and neurological disorders were also reasons for exclusion. No participant showed abnormalities in the MRI. The study was approved by the local ethics committee (Charité University Clinic, Berlin). After complete description of the study, we obtained informed written consent.

Images were collected on a 3T Magnetom Trio MRI scanner system (Siemens Medical Systems, Erlangen, Germany) using a 12-channel radiofrequency head coil. Structural images were obtained using a three-dimensional T1-weighted magnetization-prepared gradient-echo sequence (MPRAGE) based on the ADNI protocol (www.adni-info.org) (repetition time (TR) = 2500 ms; echo time (TE) = 4.77 ms; TI = 1100 ms, acquisition matrix = 256 × 256 × 192, flip angle = 7deg; 1 × 1 × 1 mm^3^ voxel size). Functional images were collected using a T2*-weighted echo planar imaging (EPI) sequence sensitive to blood oxygen level dependent (BOLD) contrast (TR = 2000 ms, TE = 30 ms, image matrix = 64 × 64, FOV = 216 mm, flip angle = 80 deg, voxel size 3 × 3 × 3 mm^3^, 36 axial slices, 5 min duration).

The first 10 volumes were discarded to allow the magnetisation to approach a dynamic equilibrium, and for the participants to get used to the scanner noise. Part of the data preprocessing, including slice timing, head motion correction (a least squares approach and a 6-parameter spatial transformation) and spatial normalization to the Montreal Neurological Institute (MNI) template (resampling voxel size of 3mm × 3mm × 3mm), were conducted using the SPM5 and Data Processing Assistant for resting-state fMRI (DPARSF, Chao-Gan and Yu-Feng [2010]). A spatial filter of 4 mm FWHM (full-width at half maximum) was used. Participants showing head motion above 3.0 mm of maximal translation (in any direction of x, y or z) and 1.0 deg of maximal rotation throughout the course of scanning would have been excluded; this was not necessary. We further analyzed head motion by correlating the frame-displacement measure (FD) with the estimated FC (see text). FD is reduced to a scalar value per each volume using the formula indicated in Power et al. [2012], and then it is averaged over volumes.

After pre-processing, linear trends were removed. Then the fMRI data were temporally bandpass filtered (0.01–0.25 Hz); but we repeated our analysis even with temporally band-pass filter (0.01 – 0.08 Hz), commonly adopted to reduce the very low-frequency drift and high-frequency respiratory and cardiac noise (Biswal et al. [1995]; Lowe et al. [1998]). The spatially normalized data were parcellated using two atlases: the automated anatomical labeling (AAL) atlas (Tzourio-Mazoyer et al. [2002]) and a recently proposed functional atlas (Shen et al. [2013]). Results for functional parcellations and for the narrow temporal filter are qualitative very similar to the ones presented in the main text, and are only reported in the supplementary material (see figures S1 and S2).

We decided to instruct participants to close their eyes during the resting state data acquisition despite the fact that resting state acquisitions with eyes open have been shown to result in slightly higher reliability of BOLD functional connectivity (Zou et al. [2015]), since the resting state data acquisition, in the longitudinal study, was part of a 1 hour scanning protocol that the participants completed every other day. Due to this fact the likelihood of falling asleep during scanning seemed particularly high to the authors and therefore the decision was taken to record all resting states with eyes closed and ask the participants after each scan session to report whether they slept during the resting state scan or not. We tested whether being asleep or not affect the distribution, but we can exclude this possibility (see figure S4 in the supplementary material). Although recently it has been recommended to acquire 10–20 min of Resting state (Birn et al. [2013]; Laumann et al. [2015]), we had to constrain data acquisition to 5 min per scan as the resting state sequence was only one of several sequences acquired in the longitudinal scan sessions. Moreover these 5 mins are representative of usual scanning times in many clinical studies.

### 2.2 Functional connectivity analysis

Spontaneous fluctuations, both at the voxel- and ROI-level, were characterized by the population variance *σ*^2^(*X*) of the BOLD-fMRI time-series *X.* For the BOLD-fMRI time-series *X = (X_1_,…, X_N_*) of a given voxel/ROI, the variance was estimated by the sample variance 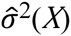

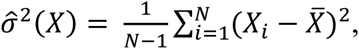

which is an unbiased estimator of *σ*(*X*). In the above equation, *X* denotes the sample mean of *X.* Functional connectivity (FC) was characterized by the population Pearson correlation coefficient *ρ*:

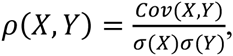

where (*X,Y*) denotes a pair of BOLD-fMRI time-series. For a pair of BOLD-fMRI time-series *X = (X_1_,…, X_N_*) and *Y = (Y*_1_*,…,Y_N_*), *ρ* was estimated by the sample Pearson correlation coefficient 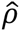:

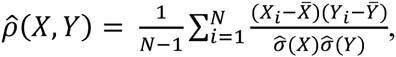

From the experimental data, we obtained, for a given subject and link, a series of sample correlation coefficients 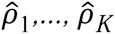, where *K* denotes the number of scan sessions. To test for non-zero inter-scan mean and variance of the corresponding population correlation coefficients *ρ*_1_,…, *ρ_K_* we used the sample mean and variance, respectively, of the series of sample correlation coefficients as test statistics. *p*-values were obtained by approximating the respective null-distributions using appropriate surrogate data (see Section 2.3) and corrected for multiple comparisons across links using the Benjamini-Hockberg method with a false-discovery-rate (FDR) of 5%.

### 2.3 Construction of surrogate data

We constructed surrogate data under the null-hypotheses of zero inter-scan FC mean and variance, based on a constrained randomization procedure first proposed in Prichard and Theiler [1994]. We first describe the construction for data from a single scan session and subsequently, describe how to use it to test for zero inter-scan FC mean and variance.

Let *X = (X_1_,…, X_N_*) and *Y = (Y_1_,…,Y _N_*) denote BOLD-fMRI time-series from two different ROI’s, where *N* denotes the length of the scan. To construct a surrogate copy of the pair of time-series (*X, Y*), the discrete Fourier transforms 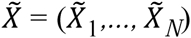 of *X* and 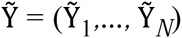 of *Y* are calculated and, subsequently, the Fourier coefficients are multiplied by random (complex-valued) phases:

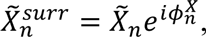

for *n = 1,…, N* and similarly for *Y.* The phases, 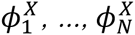 are independently drawn from the uniform distribution on the interval [0,2*π*]. Surrogate copies *X^surr^* and *Y^surr^* of *X* and *Y*, respectively, are then obtained by applying the inverse discrete Fourier transform to 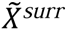 and 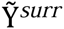.

There are two cases to consider. In the first case, the phases 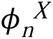 are drawn independently from the phases 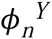, and therefore the surrogate time-series *X^surr^* and *Y ^surr^* have the same sample autocovariance functions as *X* and *Y*, respectively, but are uncorrelated. This data can hence be used to test for non-zero FC. In the second case, 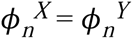, so that *X^surr^* and *Y ^surr^* have the same sample autocovariance functions as *X* and *Y* and also the same sample cross covariance function. This data can hence be used to test for dynamic FC (Hindriks et al. [2015]). We refer to these two types of surrogate data as *incoherent* and *coherent*, respectively.

To construct surrogate data under the null-hypothesis of zero inter-scan FC variance, we concatenated, for a given subject, the BOLD-fMRI data from all scan sessions, generated 1000 coherent surrogate copies, and subsequently calculated the test-statistic values to approximate their null-distribution and to calculate *p*-values.

Concatenating data from different sessions can lead to jumps in the time-series, and therefore to a possible bias in the statistical hypothesis testing. To exclude any bias, we assessed the performance of the testing procedure by generating 1000 synthetic data-sets with the same dimensions and a similar auto-correlation structure as the experimental BOLD-fMRI data, applied the procedure to test for non-zero inter-scan FC variance using *α*= 0.05, and calculated the percentage of false positives, which yielded 5.6%. When the scan sessions were shortened, the percentage of false positives remained between 5 and 6%, only increasing to 8% in the extreme case of 15 samples per scan session. This shows that the testing procedure performs well.

### 2.4 Test-retest reliability

Test-retest reliability of the functional indices was quantified by the intraclass correlation coefficient (ICC), which, for a given functional index *v*, can be defined as follows (Shrout and Fleiss [1979]). Let *v_ij_* be the measured index values of subject *i* and scan session *j*, where *i* = 1,…, *n* and *j =* 1,…, *k.*

The index is assumed to have the following form: *v_ij_ = μ + b_i_ + w_ij_*, where *μ* denotes the expectation value of *v_ij_, b_i_* denotes the random effect of the subjects, and *w_ij_* denotes all residual noise (due to dynamics, measurement error or conditions/sessions). The random variables *b_j_* and *w_ij_* are assumed to be independent and normally distributed with zero mean and variance 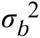 and 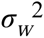, respectively. The ICC of *v* is now defined as

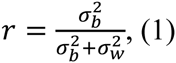

The ICC ranges between 0 and 1 and quantifies the test-retest reliability of the index *v.* Note that for an index to be reliable, it must vary between subjects (high between-subject variance 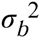) and it must be stable across scanning-sessions (low within-subject variance *σ_w_*). The most straightforward and commonly used estimator of *r*, which is sometimes referred to as the analytical estimator, is defined as 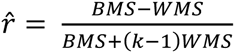, where *BMS* and *WMS* denote the mean between- and within-subjects sum of squares, respectively (here we followed the description given in Atenafu et al. [2012]). Although there are other estimators for *r*, most notably, the maximum likelihood (ML) and restricted maximum likelihood (ReML) estimators, we found them to have similar variances and only slightly different biases. The only advantage of these other estimators is the absence of negative estimates of *r*, but for simplicity we preferred to use the analytical estimator.

Statistical hypothesis testing on 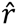 is done using an *F*-test. Specifically, since *BMS* and *WMS* are sample estimators of 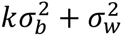 and 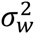, respectively, the random variable 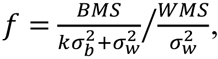, is *F*-distributed with parameters *n –* 1 and *n*(*k –* 1). Under *H*_0_, *f* takes the following form: 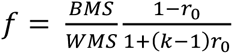, which can be used to obtain the null-distribution or 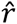.

### 2.5 Sources of variability

The issue of finite-sample variance of the Pearson correlation can be assessed by the phase-randomized surrogate data (see section 2.3). We can model the Fisher-transformed sample Pearson correlation coefficient for different participant, *i* and different scan, *j*, 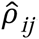, with a normally distributed variable:

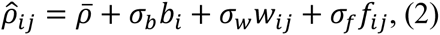

where *b, w*, and *f* are independent, and standard-normally distributed random variables. The random variable *w* models the genuine variability of FC in each subject (within-subject), the random variable *b* models the FC variability for different subject (between-subject) and the variable *f* models the finite-sample error. Assuming *σ_w_* to be independent of *i* (subject) means that the genuine variability of FC over scans, as measured by *σ_w_*, is equal for all subjects (a strong assumption).

The three sources of variability can then be separated and the three variances, 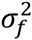, 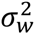, and 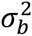 can be calculated from the surrogate analysis:

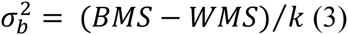

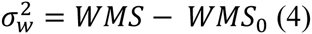

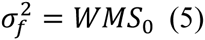

where *WMS* and *BMS* are the mean square errors within and between subjects, *WMS*_0_ is the mean square error within subjects for the surrogate case, and *k* is the number of sessions. From this we get the ICC value:

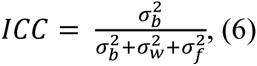

Since the surrogate data is constructed under the null hypothesis of no genuine FC variability over scans (that is, *σ_w_* = 0), the ICC constructed from the surrogate data, ICC_0_ equals

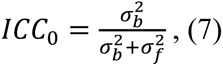

and therefore, ICC_0_ ≥ ICC, so that the surrogate data can be used to estimate the uncertainty in the ICC that is due to the finite-sample size.

### 2.6 Definition and estimation of functional similarity

Central to the analysis in Finn et al. [2015] (but see also Mueller et al. [2013]) are the within-and between-subject *similarity indices*, here denoted by 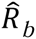 and 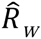, respectively. *R_w_* can be calculated for every subject *i* and for every pair of scan sessions (*j, j*′) and is defined as the sample Pearson correlation coefficient between the respective vectorized (and *z*-scored) FC matrices *X_ij_* and *X_ij_*_′_ with *j ≠ j*′:

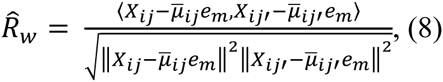

where *μ_ij_* is the average over the links of *X_ij_*, 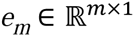 denotes the vector containing all ones and where we have suppressed the dependence of 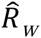 on (*i, j*′), and *j*′ from the notation. Similarly, 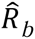 can be calculated for every two subjects *i* and *i*′ (*i ≠ i*′:) and every pair of scan sessions:

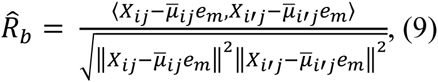

Note that *R_w_* and *R_b_* can be used to assess the similarity not only for the vectorized FC matrix, but for *any* multivariate biomarker. In the sequel, therefore, we let *X_ij_* denote an arbitrary *m*-dimensional biomarker for subject *i* and scan session *j*.

To assess the properties of 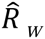 and 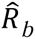, we need to consider the respective population quantities, which we will denote by *R_w_* and *R_b_*, respectively. Below, we denote *X_ij_* for the observed value of the biomarker and *x_ij_* for the corresponding population biomarker (*X_ij_* is a *realization* of *x_ij_*). The definitions of *R_w_* and *R_b_* are obtained by replacing the sample Pearson correlation coefficients in Equations (8) and (9) by the population Pearson correlation coefficients and replacing *X_ij_* by *x_ij_*:

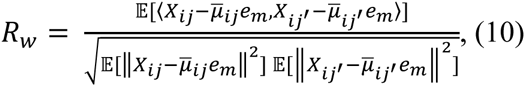

for *j ≠ j*′ and

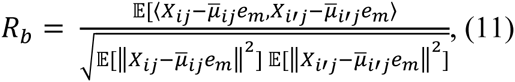

for *i ≠ i*′. To assess the properties (bias and uncertainty) of the estimators 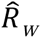 and 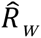, we also need a statistical model for the population biomarker *x_ij_*. This will be described in the next section.

### 2.7 Statistical model for multivariate Gaussian biomarkers

Let 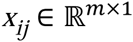 denote an arbitrary *m*-dimensional (population) biomarker of subject *i* (*i* = 1*,…,n*) on scan session *j* (*j* = 1*,…,k*). In analogy to the univariate linear model used to assess local test-retest reliability, we model *x_ij_* by the following multivariate linear model:

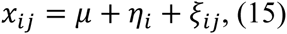

where 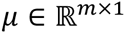 denotes the group-wise expectation of *x_ij_*, and where 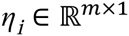 and 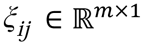 denote within- and between-subject fluctuations, respectively. The random vectors *η_i_* and *ξ_ij_* are assumed to be independent and have expectation zero (that is, the *m*-dimensional zero-vector) and covariance matrices Σ*_w_* and Σ*_b_*, respectively. Note, that Σ*_w_* and Σ*_b_* are the generalizations to the multivariate case of the within- and between-subject variances 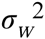 and 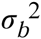, respectively.

Assuming in first approximation that Σ*_b_* and Σ*_w_* are diagonal matrices, the expectations of the similarity indices *R_w_* and *R_b_* can be expressed in terms of the model parameters as

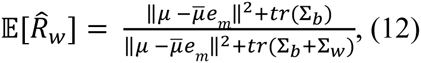

and

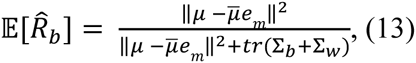

where ***tr*** denotes matrix trace and 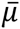 denotes the average value of *μ*. As a special case, suppose that Σ*_b_* and Σ*_w_* are identity matrices multiplied by a factor, that is 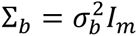 and 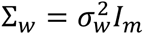 for certain *σ_b_* and *σ_w_* Then Equations 12 and 13 reduce to

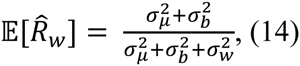

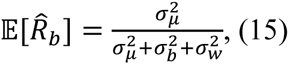

where we defined 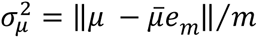.

It is possible to derive approximate formulas for the expectation of the similarity indices, for the more general case:

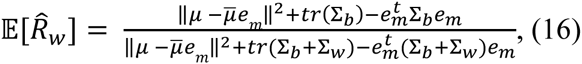

and

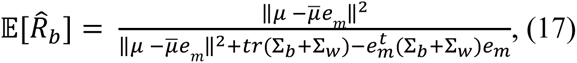

The variances of the similarity indices were approximated using Equation 3.1 in Dutilleul et al. [1993]:

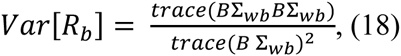

and

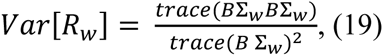

where Σ*_wb_*, and Σ*_w_* _+_ Σ*_b_*, *B* = (*I_m_* – *J_m_* / *m*)/*m* and *I_m_* and *J_m_* denote *m*-by-*m* identity matrix and the matrix of ones, respectively.

We checked the feasibility of this approximated formula using simulated data and found that it is an upper bound for the indices. In the simulations, we generated synthetic connectivity matrices *FC_ij_ (i* subjects, *j* sessions), with a multivariate general linear model, and using for matrices 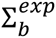 and 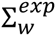 the estimated values obtained from the data. To simulate different conditions, as the ones analyzed in the experimental data set, we fixed Σ*_b_* for several simulations and used different matrices 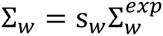, where the parameter s*_w_* was used as a multiplicative factor.

## 3 Results

We present a systematic analysis of the variability and reliability of resting-state functional connectivity, both at the level of individual ROI pairs and of the entire brain. We used five-minute resting-state fMRI data from six participants, each of which was scanned 42 times (see Material and Methods for detailed information on subjects and pre-processing). ROI-level analyses were conducted using an anatomic parcellation (AAL, Tzourio-Mazoyer et al. [2002]) and the main results were replicated using a functional parcellation (Shen et al. [2013], see supplemental information).

### 3.1 Data set description

Before moving into the details of the analysis, we want to give a descriptive overview of the data-set to provide the reader with an intuition for how variable and reliable functional connectivity is. As a first step, we look at the inter-session variability of the average FC over links, <FC> (see panel A of figure 1). For the 5 subjects scanned multiple times, the average interval of time between the first and the last session spanned approximately 6 months. It is possible to appreciate that in general the average <FC> for the 5 subjects scanned multiple times (blue dots) resembles that computed on the 50 subjects, each of which scanned just once (gray continuous line). The same effect can be observed in panel B, in which we can compare the distribution of <FC> for the 5 subjects (blue lines) and the distribution of <FC> for the 50 subjects (gray bar).

**Figure 1:**
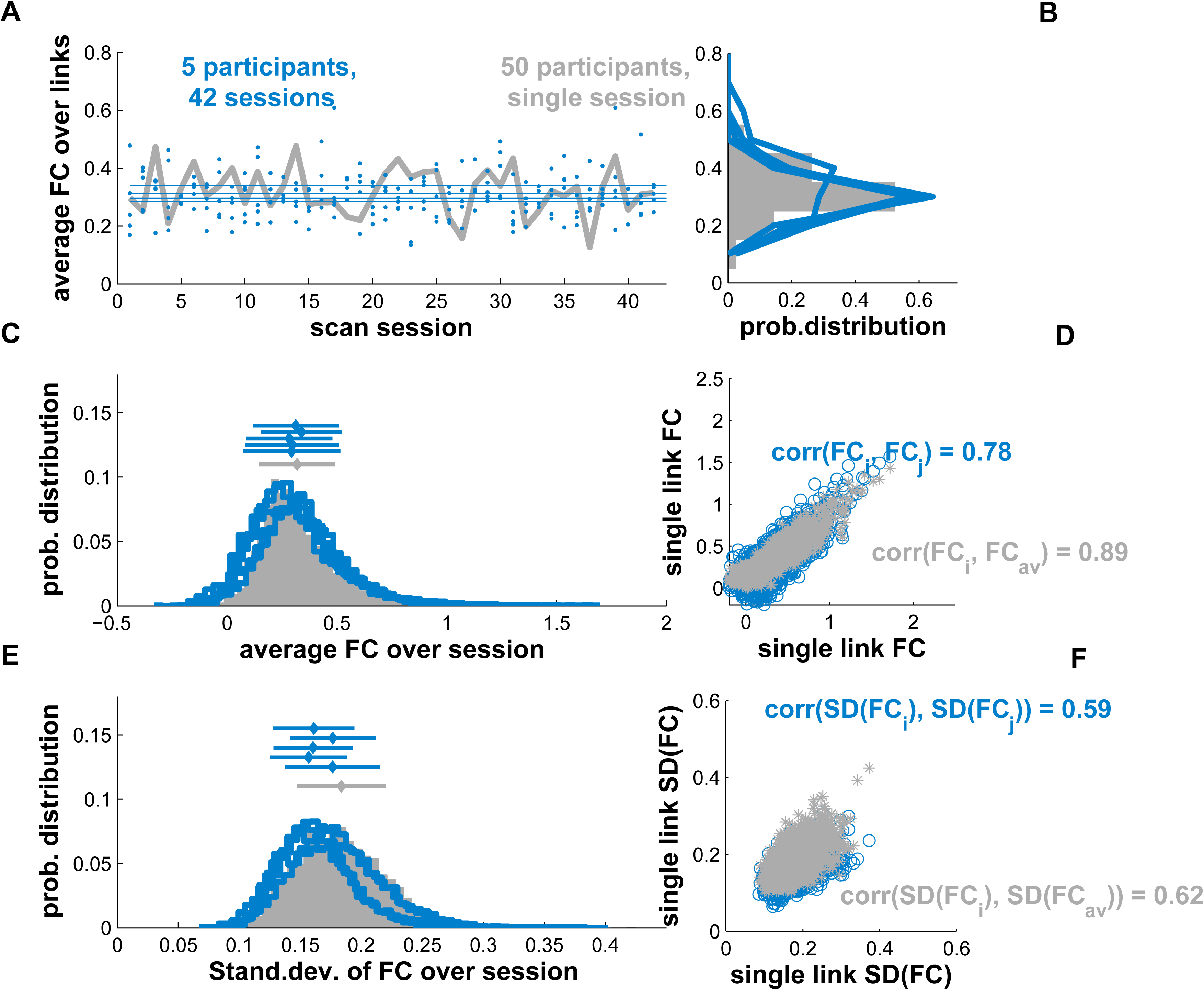
Variability of FC. Panel A: Average FC over links, <FC>, of the 5 subjects for all 42 sessions (blue dots), and for the 50 subjects (gray line). Panel B: distribution of <FC> of the 5 subjects (blue lines) and the distribution of <FC> for the 50 subjects (gray bar). Panel C: the distributions of the FC values (blue lines for the 5 subjects, and gray bar for the 50 subjects). Panel D: the FC values of a participant (FC*_i_*) against FC values of another participant (FC*_j_*, blue circles), and against the FC values of the average of the 50 subjects (FC_50_*_s_*, gray asterisks). In the same panel, we reported the correlation between two participants’ FC (corr(FC*_i_*,FC*_j_*) ≈0.8), and the correlation between a subject’s FC and FC_50_*_s_* (corr(FC*_i_*,FC*_j_*) ≈0.87). Panel E: the distributions of the standard deviation over sessions of the FC (SD*_FC_*). Same color conventions as panel C. Panel F: one participant’s SD*_FC_* against the SD*_FC_* of another participant (blue circles), and against the SD*_FC_* of the 50 subjects (gray asterisks). In this panel, we reported the correlation between two participants’ SD*_FC_* (averaged over 42 sessions), and the SD*_FC_* of a subject against the ones of the 50 subjects.

In panel C of figure 1 we can see how the distributions of the FC values for the 5 subjects scanned multiple times (FC*_i_*, blue lines), and of the FC of the 50 subjects (FC_50_*_s_*, gray bar) are very similar, with a similar average; The distributions of all FC values for the 5 subjects and that of the 50 subjects are in general very similar, even though the latter is narrower with a standard deviation (SD) of 0.35 compared to the former, whose SD is 0.45. Another way of measuring the similarity between FC_50_*_s_* and FC*_i_* is through their correlation. In our data set, the average correlation between any couple of FC*_i_* is 0.8 (SD=0.02), and the correlation between an FC*_i_* and FC_50_*_s_* is slightly higher, 0.87 (SD=0.02, see the scatter-plot of panel D). Therefore, the average FC_50_ for the 50 subjects scanned just once can be considered as representative of the FC obtained from single individuals. To complete this preliminary description, we look at the inter-session variability of the 5 subjects’ FC and subject-by-subject variability of the FC_50_ (panel E and F). We note the high similarity between the distributions of the standard deviation over sessions of FC*_i_* (SD*_FCi_*) and of FC_50_*_s_* (SD*_FC_*_50_*_s_*). However, we observed rather low values of correlation between the SD*_FCi_* of any two of the 5 subjects (0.53, SD=0.02), indicating high variability between subjects of spatial distribution are quite different from subject to subject, and variability between-subject is even less representative.

### 3.2 Link-wise analysis

#### 3.2.1 Within-subject variability

We now present the analysis of reliability and variability of single links. Within-subject mean and variability of a given link’s correlation were quantified, respectively, by the sample mean and standard deviation, SD, of the corresponding time-series of correlation coefficients. By repeating the calculations for each link, two matrices for each subject and for the 50 subjects were obtained, corresponding to the within-subject average FC and variability (standard deviation, SD, of FC). In Figure 2, we show these matrices with heat-maps for the 50 subjects (panels A-C, blue), and for one of the subjects (panels B-D, cyan). To represent a more robust measure, we averaged the links over macro-regions (see labels in the panels). The ROIs for this figure were defined with AAL parcellation (see supplementary material for the corresponding plots with Shen’s parcellation, with very similar results). They show that both the average FC and standard deviation vary considerably over links. Note also the existence of relatively high SD values for some links. This suggests that the FC strength between corresponding ROI’s varies considerably from scan to scan.

**Figure 2:**
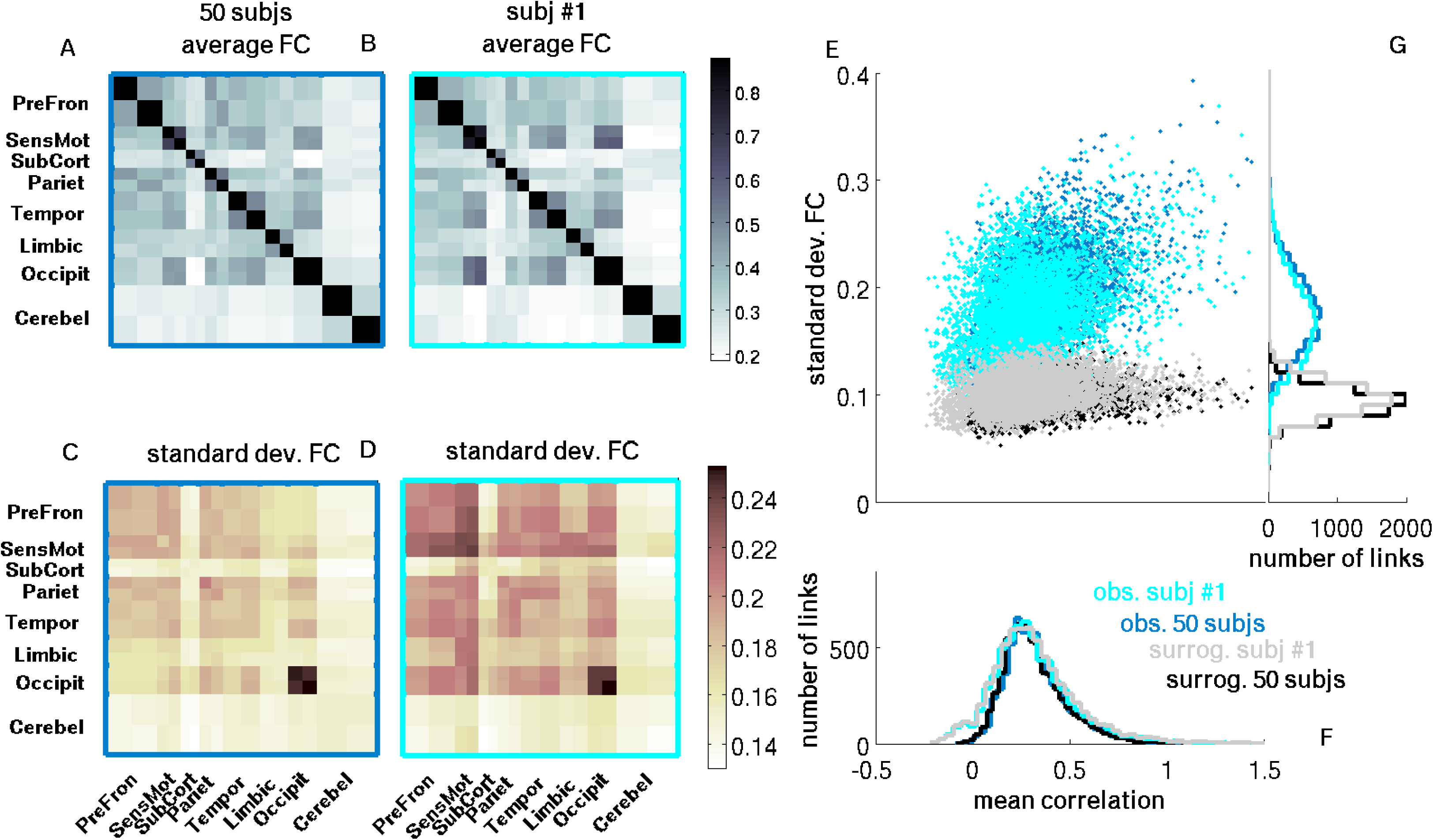
Between- and within-subject FC variability. Panels A-B show the heat-maps of the average FC for a single subject and for 50 subjects, respectively. Panels C-D show the FC standard deviation (SD) for a single subject and for 50 subjects, respectively. The average FC is Fisher-transformed (inverse hyperbolic tangent), and the SD is calculated from these transformed values. Color convention is cyan and blue indicate 50 subjects and single subject, respectively; black and gray dots indicate surrogate data for the 50 subjects and the single subject, respectively. Panel E shows the scatter-plot of the average FC against the FC standard deviation. Panels F and G plot the distributions for average FC and SD with the same color conventions. All the plots of this figure refer to one exemplary participant. The figures for the other four participants are qualitatively similar, but not reported.

Panels E-G display the same average FC and its standard deviation with scatter-plot and histograms. Average and standard deviation are calculated over scans (this means over sessions for the single subject and over subjects for the 50 subjects). For both single subjects with multiple scans (cyan) and 50 subjects (blue), FC’s standard deviation ranges between 0.1 and 0.3, with an average value of about ≈0.2 and a SD of about ≈0.038 (for the single subject with multiple scans the range the SD is slightly inferior 0.035). We can see that the value of the average FC is a factor influencing the variability of the FC itself: the correlation between average FC and its SD is ≈0.56 (≈0.4 for the single subject).

##### Testing the null-hypothesis of static FC

Does the observed variability of the FC reflect genuine variability of spontaneous inter-areal co-activations, or does it arise from mere statistical uncertainty of estimates? It should be remembered that Pearson correlation coefficients are sample estimates of the population values (see Materials and Methods), and as such, finite-sample variability should not be confounded with the variability due to real underlying dynamics of the FC (see for example Lindquist et al. [2014]; Hindriks et al. [2015]).

With this in mind, we tested the null-hypothesis that the observed fluctuations of FC can be fully explained by statistical uncertainty of the correlation estimates: to this aim, we first constructed appropriately randomized data Prichard and Theiler [1994] (see also Materials and methods). This randomization method yields data with the same statistical structure and the same mean FC as the empirical data, but introduce no dynamics (see Materials and Methods for more details); from now on, we will refer to these randomized data as surrogates. In panels E-G of Figure 2, one realization of the surrogates is plotted (black and gray lines and circles). From see panel G, it is evident that the distribution of the SD of the correlation for the surrogates (black and gray) are qualitatively different from the observed data (blue and cyan). By construction, the distributions of the mean correlation of the surrogates (blue) and of observed data (black) are identical (panel D). Using this method, and by repeatedly randomizing the data multiple times, we can approximate the distribution of the variability for each functional connection under the null-hypothesis of constant FC variability of each link, and hence *p*-values can be calculated. Applying the Benjamini-Hochberg method for multiple comparisons with a false-discovery rate (FDR) of 5% we found that the approximate number of functional links whose variability can be explained by the null-hypothesis of no genuine variability is around 1%. This means that, for each of the five participants, practically every functional connection varies over different scanning sessions: in other words, the day-by-day co-variation between different brain regions appears to be dynamic.

#### 3.2.2 Test-retest reliability

In statistics, the reliability of a measure indicates its consistency under similar conditions in contrast to dissimilar conditions: therefore, a measure is highly reliable and amenable to be a good biomarker, if it yields similar results under consistent conditions, but not under dissimilar conditions. An example of a reliable measure is people’s height, which tends to be stable for a given individual, but exhibit large variability across individuals. How reliable are the pairwise functional indices obtained in typical resting-state studies? To address this question, we measured how stable the FC estimates of the same participant were over different scan sessions (within-subject variability) compared to those obtained from different participants (between-subject variability). For the functional connectivity estimates to be considered reliable, according to the definition of reliability mentioned above, they should therefore exhibit small within-subject variability while at the same time large between-subject variability.

Following previous studies (Shehzad et al. [2009]; Zuo and Xing [2014]), test-retest reliability of the functional indices was quantified by the intraclass correlation coefficient, ICC (see details in Materials and Methods). Estimated ICC’s for all links are plotted in panel A of Figure 3 against the subject-averaged FC (gray asterisks), while panel B shows the histogram of the estimated ICC values. In the legend we indicated the gray histogram as ‘observed’ in contrast to the values obtained with the simulation and the theoretical analysis (see below). The heat-map in panel C shows ICC values averaged over the regions indicated in the labels.

**Figure 3:**
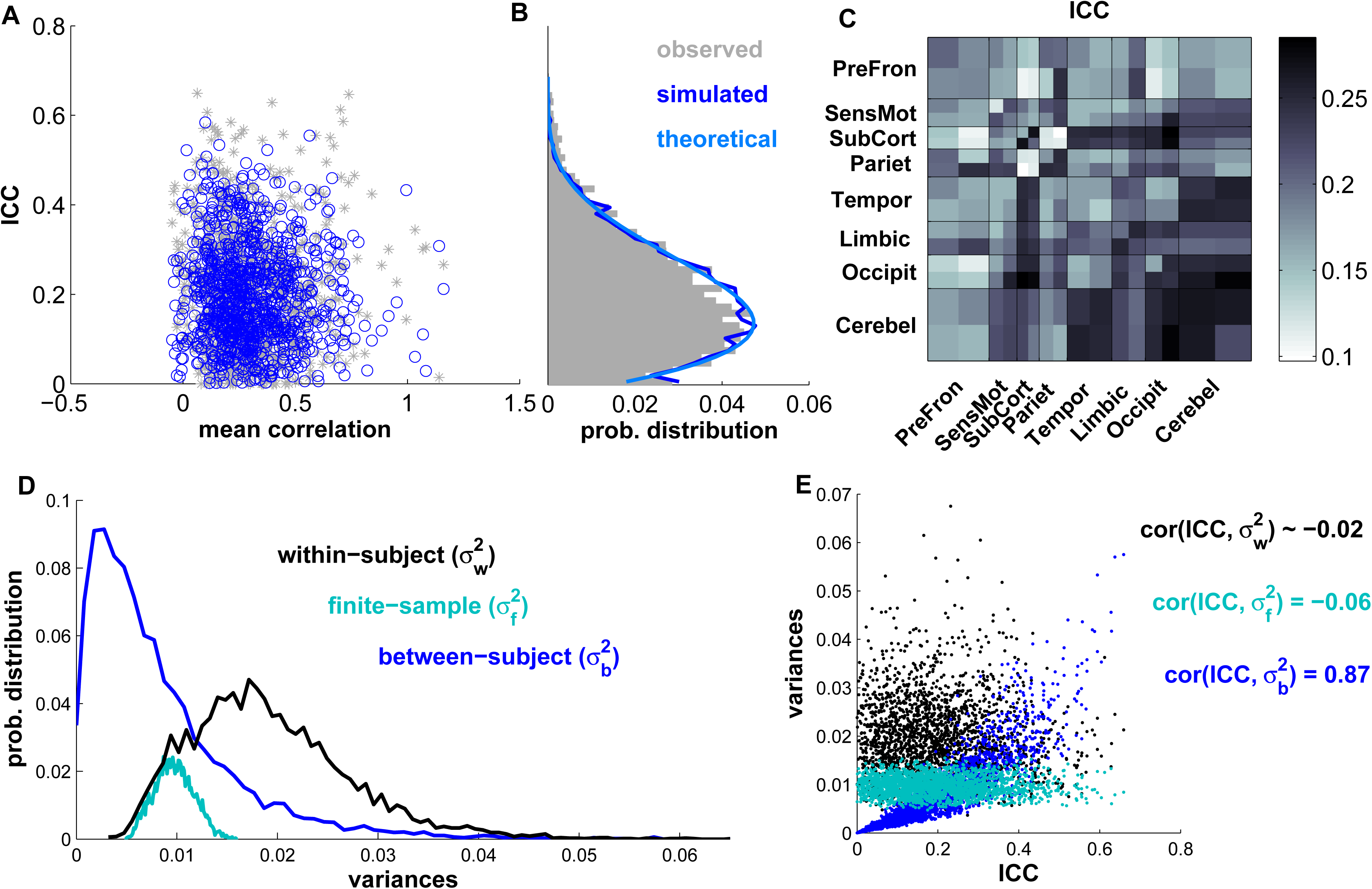
Reliability of the correlation strength. Panel A shows the scatter-plot for the correlation strength against ICC value, panel B shows the histogram for the distribution of the ICC values, and panel C is the heat-map of the average ICC values for the different macro-regions. For the panels on the left and on the center, the colors light blue, dark blue and gray refer to the theoretical, simulated and observed values, respectively (see main text). Panel D shows the distribution of the three variances (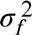, 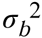, and 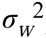). Panel E shows the scatter plot of the three variances against the ICC, with the values of the correlations between the three variances and the ICC; the colors follow the same convention of panel D.

Note that the estimated ICC values vary from link to link, ranging from approximately 0 to about 0.7. With the estimator we used, negative values of ICC can be obtained. Estimators of ICC based on the likelihood estimation can circumvent this problem, but as the two estimators showed no qualitative differences, we used the simple estimator. The average value of the ICC is ≈0.22 ±0.16, which is commonly considered rather low and indicates that link-wise FC for 5 mins scan performs poorly as a biomarker for individual subjects Nunnally [1994]. Despite the current absence of common consensus about what should be considered an acceptable level of reliability (Nunnally [1994]; Lance et al. [2006]), it is not debatable that ICC around 0.2 is a poor value. Indeed to observe such low ICCs, the within-subject variance has to be twice as large as the between-subject variance. Such a low ICC mirrors the fact that the within-subject variance is twice as large as the between-subject variance: low within-subject variability compared to between-subject variability.

Earlier studies have reported similar values for the ICC Shehzad et al. [2009]; Zuo and Xing [2014], but with substantial differences in their interpretation (see below). Similar results have been observed also by Birn et al. [2013], even though these data is more difficult as they were obtained by combining different sessions, and with three different conditions (eyes-open, eyes-closed and fixation).

The variability of ICC estimates across links is also in line with previous reports with similar duration scans Shehzad et al. [2009]; Zuo and Xing [2014]. In those studies, this variation was interpreted as evidence for heterogeneity of test-retest link-wise reliability of functional connectivity (among other biomarkers), but the statistical variability of the ICC estimates was not explicitly considered. It remains therefore possible that the observed variability in ICC values reflects statistical uncertainty, rather than true variation of the ICC across links. Indeed, even with 42 scan sessions and 5 subjects, the variance of the ICC estimators is considerable. We therefore tested the null-hypothesis of all links having the same population ICC. The population ICC*_av_* under the null-hypothesis was thus considered to be the estimated link-wise average ICC.

We first calculated the probability of each link to have such a value of ICC, or higher, given the assumption of being an estimate of ICC*_av_*. Thus, this probability corresponds to a p-*value*. Then we calculated how many links had an ICC statistically different from ICC*_av_* after a false discovery rate correction (using Benjamini-Hochberg method with FDR=5%).

In panel B of figure 3 the distribution of observed ICC (gray bars) and the theoretical distribution of ICC (light blue line) can be found, estimated from a general linear model (GLM) with a constant theoretical ICC (see section 2.4 for details). As mentioned above, the average theoretical ICC value chosen was ICC*_av_*. We can see that the three distributions are practically identical, demonstrating that there is no link having an ICC different from ICC*_av_*. The used statistical framework suggests that, as a first approximation, the ICC of each individual link can be drawn from a unique distribution, hence proving strong evidence that FC test-retest reliability is homogeneous over links.

To further test this hypothesis, we simulated the links correlation variability with Gaussian stochastic variables having two sources of variability, ‘within-subject’ and ‘between-subject’. Each simulated correlation was generated as a Gaussian variable, whose average value is equal to the observed mean correlation of one real link: 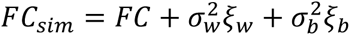; where the variances, 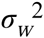 and 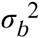, were maintained constant for all the simulated correlations. The ratio between the two variances was chosen equal to the average ICC*_av_*, and for simplicity we set 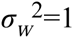 (the actual value does not influence the results of the simulation). We extracted these variables one time per each simulated subject, and 50 times per each simulated scan session. Finally, we calculated the ICC values for each simulated correlation, *FC_sim_*. The results of this simulation are reported in panel A as blue circles, and their distribution in panel B with a dark blue line. Even in this case, it is possible to appreciate that the simulated distributions very well approximates the empirical one.

It should be stressed that one possible explanation for the lack of heterogeneity in the ICC values of the links might be the lack of statistical power: few subjects, limited number of scan sessions, correction for multiple comparisons, etc. To circumvent the issue of having to perform a too restrictive multiple-comparison correction, we took the average ICC’s of different macro-regions (the names of these regions are indicated in the labels). Here, we use the term macro-region to indicate a brain region composed of several ROIs. The idea is based on the hypothesis that different macro-regions might have different reliabilities. This approach closely follows that taken in Zuo and Xing [2014] in which systematic differences were reported of ICC’s averaged over the different resting-state networks (RNSs) for several functional biomarkers, including (intrinsic) FC.

The heat-map in panel C of figure 3 shows a synthetic picture of ICC of the average correlation between pairs of macro-regions. The differences are small: all the average ICCs range between 0.1 and 0.3. We compared the ICC distribution between pairs of macro-regions with a non-parametric test (see Methods for details), and we did find most of them to be statistically different. Therefore we can sort the macro-regions according to average ICC value, and isolate the least reliable region (parietal region, whose average ICC≈0.15) and the most reliable one (cerebellum, whose average ICC≈0.24). The least reliable macro region outside itself is the pre-frontal one (whose average ICC=0.18) and the most reliable one is cerebellum, whose average ICC=0.27.

#### 3.2.3 Sources of variability

We now analyze the different sources of variability of the FC, and how they relate to the ICC reliability. We can indeed disentangle the contribution of three different sources of variability: 1. the genuine variability of FC in each subject, within-subject variability; 2. the FC variability for different subjects, between-subject variability; 3. the variability due to the statistical uncertainty associated with computing the correlation from a finite number of samples, finite-sample variability. We note that, while the first two sources have already been partially accounted for in the literature, the finite-sample variability has not been explicitly addressed before (Shehzad et al. [2009]; Birn et al. [2013]; Zuo and Xing [2014]; Laumann et al. [2015]). Note that the description in terms of these variability sources is slightly different from what has been reported in literature (see i.e., Zuo and Xing [2014]; Laumann et al. [2015]; Mueller et al. [2013]). For example, 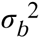, defined as the between-subject variance is not obtained calculating the variances between the sessions of different subjects, that instead should be approximated by the sum of 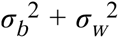 (Laumann et al. [2015]). As we are describing the correlations as Gaussian variables, each source of variability is associated with a corresponding variance: between-subject variance 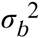, finite-sample variance 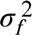, and within-subject variance 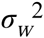.

Note that for each subject, the inter-session variability of the correlations can be divided into within-subject variability and finite-sample variability. To calculate the contribution of the finite-sample variability, we used the surrogate data described before, as they possess finite-sample variability, but not, by construction, within-subject variability (see Materials and Methods). Therefore, to obtain 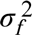, for each link, we subtracted the value of the inter-session variability obtained from the observed data to the one obtained for the surrogate data. For the observed data, both the finite-sample variability and the between-subject variability resulted on average approximately half of the within-subject variability; see the complete distribution of the three variances in panel D of figure 3. This strong difference between 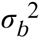 and 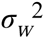, evident from their distributions, is the main reason for the low reliability of the links, described in the previous section.

The values of the three variances averaged over regions are reported in figure S5 of the supplementary material. Although the variances present homogeneous values for all the regions and it is not clear a pattern, we note that both the ICC and the three variances form a characteristic structure, with some macro-regions exhibiting different patterns of behavior with respect to the others (see e.g., the occipital).

We also analyze how the three variances correlate with the ICC (see panel E of figure 3: the correlation of the ICC with the 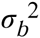 is very high (≈0.86), while the correlations with other two variances is almost zero, 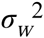 (-0.07, *p*-value > 0.05) and 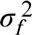 (-0.05, *p*-values < 10^−5^). These results have a straightforward interpretation: the between-subject variability represents a structure similar to the one of the ICC, while the differences between the regions in the within-subject variability are not strongly related to the region differences in the ICC.

Recently, different studies have warned against the influence of head-motion and micro-movements (i.e. head displacements <1mm), in the observed variability and reliability of FC estimates (Power et al. [2012]; Laumann et al. [2016]). Taking into account this possibility is indeed very relevant for our analyses, as it indicates one of the different plausible causes behind within-subject variability, namely unavoidable head movements during the scan session, which should in principle being independent from scan to scan. To assess this possibility, we calculate the correlation between the inter-session variability of the average frame-displacement (FD) and that of each link’s correlation (see Methods for the calculus of FD). We found that head-motion explains part of the variance of the correlation (≈ 5%), even though the effect is not homogeneous over different regions, see panel A of figure S5 of supplementary material. Moreover, the FD-effect correlates positively with within-subject variability (≈ 0.35), but not with between-subject variability.

#### 3.2.4 Relevance of sample points: scan duration and multiple scans

Different studies have analyzed the effect of scan duration on the reproducibility of FC (Anderson et al. [2011]; Birn et al. [2013]; Hacker et al. [2013]; Laumann et al. [2015]; Finn et al. [2015]), and on the reliability (Shehzad et al. [2009]; Birn et al. [2013]) demonstrating that long scan sessions increased both the reliability of FC and its reproducibility. We note that the former is not an obvious consequence of the latter, in that having highly reproducible FC within-subject could also mean highly reproducible FC between-subject and therefore low reliability. For example, Birn and colleagues demonstrated that reliability slowly increased with scan duration: on average, the maximal ICC value for very long scans (30 mins) is very low, ICC≈ 0.4 (Birn et al. [2013]).

As such, we systematically studied the influence of scan duration on the reliability of FC indices. Moreover, we analyzed the behavior of the ICC as a function of the different sources of variability (within-subject, between-subject and finite-sample). Panel A of figure 4 shows the behavior of the three variances and that of the ICC for different scan durations, as quantified by minutes.

**Figure 4:**
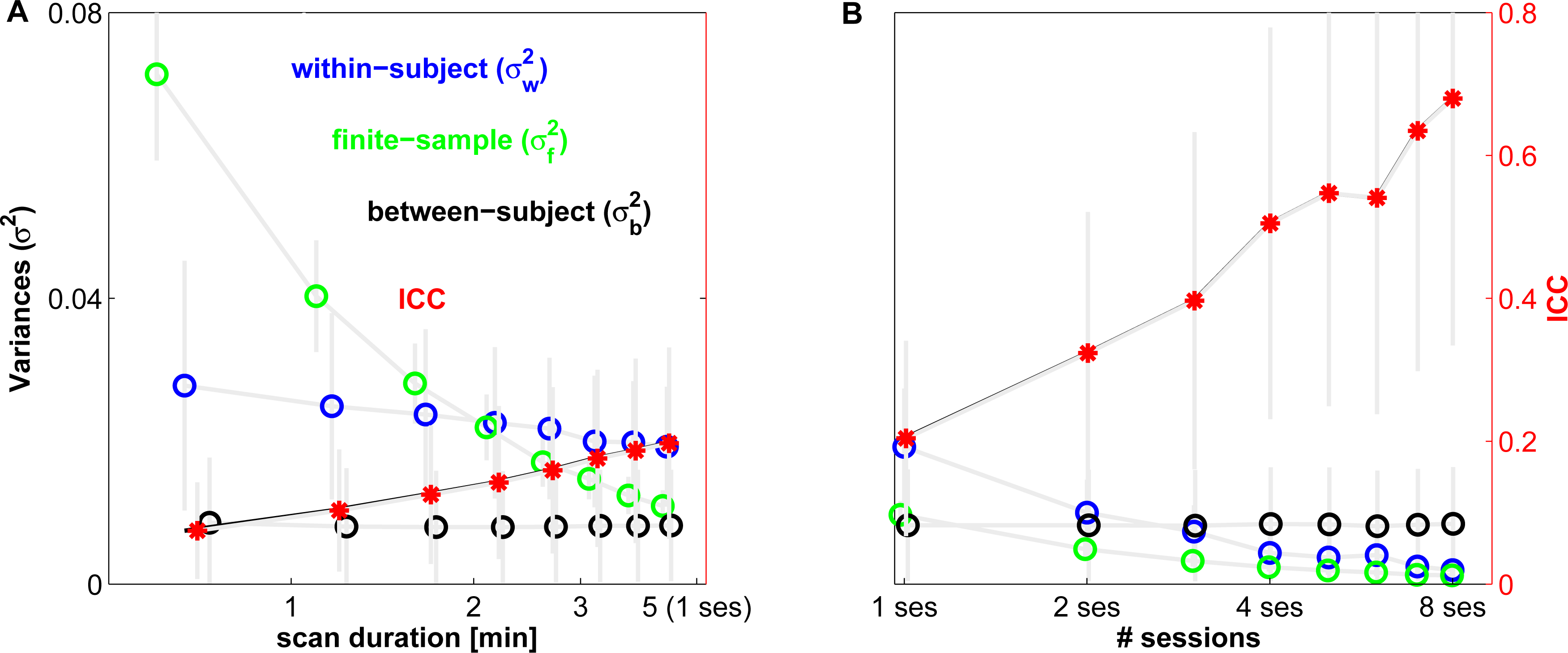
Effect of scan duration on the FC reliability. The graph shows the behavior of the average reliability, ICC, and the behavior of the three variances related to the three sources of variability of FC for different scan duration (panel A) and using multiple scan sessions (panel B). The empty circles refer to the three sources of variability: within-subject (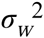, blue), finite-sample (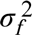, green) and between-subject (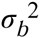, black). The red asterisks refer to ICC. To plot ICC we used a second y-axis (in red, on the right). In gray the SD of each measure are reported. The points are slightly misaligned to improve the plot readability.

For very small scan duration (below 1 min), finite-sample, 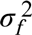 (green line) is the most relevant source of variability of the FC indices, even though its contribution rapidly decreases with increasing scan duration. We observed that the behavior of finite-sample variability can be approximated by a power law of 1/*N^a^*, where N is the number of time points, and *a* is about 1.3 (*χ*^2^ = 0.98). This is not surprising, as the finite-sample variance of the correlation between any two time-series having zero auto-correlation is equal to one. Within-subject variability (blue line) also tends to decrease with increasing scan duration, even though at a much slower rate, whereas between-subject variability (black line) remains approximately constant.

Having many scan repetitions obtained from the same subject, we could also measure both the three variances and the ICC obtained using the average FC over several sessions (details can be found in the Materials and Methods). Results from this analysis are depicted in panel B of figure 4, in order to directly compare them to the evolution of the variances for different scan duration. We note that the between-subject variability 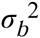 again tends to remain constant, whereas the finite-sample variability 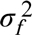 continues to decrease with no evident changes in slope. On the other hand, within-subject variability 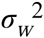 seems to exhibit discontinuous changes that are mirrored by abrupt changes in the slope of ICC. These abrupt changes are expected given the previous results reported in literature (Shehzad et al. [2009]; Birn et al. [2013]) on the difference between the reliability within-scan session (less than one hour) and between-scan sessions (more than one month), with higher values of reliability for the case within-scan session. This result indicates that to obtain intermediate or high level of reliability, we should average FC over multiple sessions. Indeed, according to the results of Birn et al. [2013], there seems to be a plateau for the ICC between-scan sessions above 18 minutes (see figure 3a of Birn et al. [2013]). Evidence for this slope change can be found in the high value of ICC (≈0.7) obtained for FCs extracted from an average of 6 sessions (summing up to approximately 30 mins).

We underline the relevance of this analysis: First, we can describe how reliability of the FC changes as a function of scan durations or using several scan sessions for the three different types of variance, and second, we conclude that the use of multiple sessions seems to be a potential way to overcome the low reliability upper limit indicated by Birn et al. [2013]).

The relevance of the finite-sample variability is conspicuous, but it will shade out for increasing scan duration. The influence of scan duration on the sources of variability will be the subject of the next section.

### 3.3 Global FC analysis

After having analyzed reliability from a local, link-wise perspective (Section 3.2), we focused on studying the inter-scan variability of the whole-brain, global FC structure. This means that instead of considering the variability of the different pair-wise functional correlations, we consider the within- and between-subject variability of the vectorized FC matrices in their entirety. As in the local analysis (see Section 3.2), and following earlier studies Mueller et al. [2013]; Laumann et al. [2015]; Finn et al. [2015], functional connectivity was quantified using Pearson correlation coefficient. The richness of information contained in the multivariate structure of whole-brain resting-state FC matrices has recently been demonstrated Finn et al. [2015]. In that study, it was shown that the FC matrix can be used as a “functional fingerprint” in that it allows identification of individual subjects from a 30-minute resting-state scan. The findings in Finn et al. [2015] appeared to be in stark contrast with the low test-retest reliability of local FC indices. For example, Birn et al. [2013] reported low ICC’s (≤ 0.4) for pair-wise Pearson correlations even for long scan sessions (30 min) and we reported similar values (see Section 3.2).

In this section we reproduce the findings in Finn et al. [2015] (Section 3.3.1), providing a statistical framework that can be used to assess the factors influencing functional fingerprinting (Section 3.3.2). Taken together, our results confirm the strength of whole-brain FC analysis over local measures.

#### 3.3.1 Subject identification from resting-state FC

In this section, we reproduce the observations of Finn et al. [2015] and again assess the effect of scanning duration. The analysis carried out in Finn et al. [2015] is based on the sample Pearson correlation coefficients between different pairs of vectorized FC matrices, to which they referred to as *similarity indices*. These similarity indices can be calculated between (vectorized) FC matrices of different scans of the same subject (within-subject) or between FC matrices obtained from different subjects (between-subject). The within- and between-subject similarity indices are denoted here by *R_w_* and *R_b_*, respectively. Details are provided in Section 2.7. Finn and colleagues demonstrated that for 30-minute resting-state scans, *R_w_ > R_b_*, for practically all values of *R_w_* and *R_b_* (calculated from all possible pairs of scan), which implies that *R_w_* and *R_b_* can be used as “functional fingerprints” to identify individual subjects. We repeat the analysis of F2015, calculating the distribution of *R_w_* and *R_b_*, collapsing together different sessions. In figure 5, panel A,C-E show the observed distributions of (Fisher transformed) *R_w_* (gray) and *R_b_* (black) for a different numbers of samples (number of sessions). It is possible to appreciate that the separation between the distributions of the two similarity indices, *R_w_* and *R_b_*, increases rapidly when increasing the number of sessions: from panel A (1 session) to panel E (6 sessions). This separation is almost complete (zero overlap between the distributions) even with 4 sessions. This is noteworthy as with 4 sessions the average ICC of single link is still around 0.4 (similar value is reported by Birn et al. [2013]), and however on the whole-brain level they allow for complete identifiable FC (Finn et al. [2015]).

**Figure 5:**
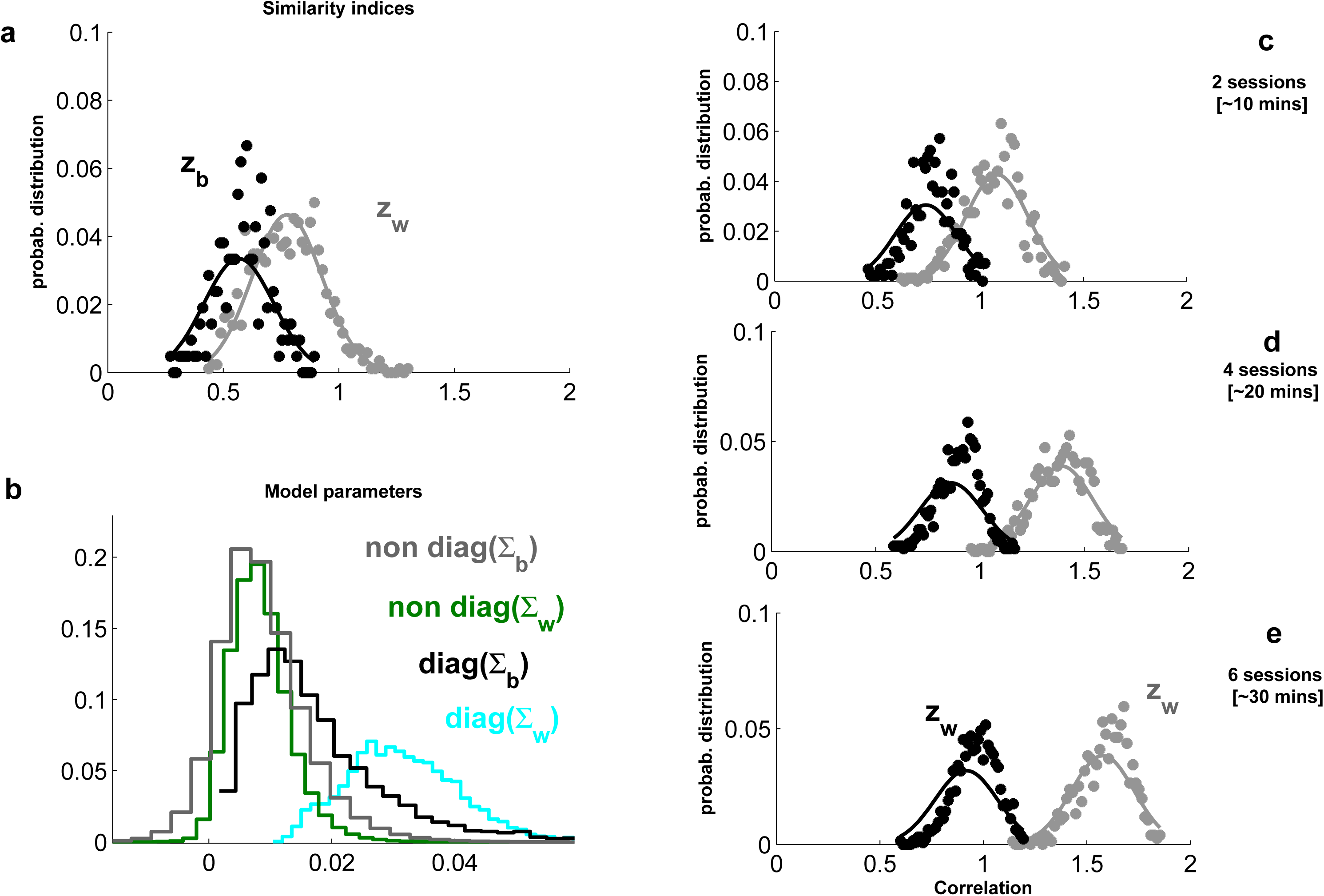
Analysis of FC at global level. The two panels on the left refer to the analysis done using the single session. Panel B shows the distributions of the estimated parameters’ values with the general linear model. Panels A, C-E plot the distribution of *z_w_* (gray) and *z_b_* (black) for FC averaged over 1, 2, 4, and 6 sessions, respectively. The observed data are represented with dots, and the theoretical approximated values with continuous lines. The separation between the distributions of *z_w_* and *z_b_* increases rapidly when increasing the number of sessions.

To explain why functional fingerprinting is possible and how its quality depends on different factors (scanning length, for example), we constructed a statistical model for the vectorized (and *z*-transformed) FC matrices. Specifically, the vectorized FC matrix of subject *i* at scan *j*, denoted by *x_ij_* is modeled as a normally distributed random vector having the following structure:

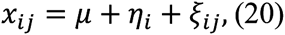

where 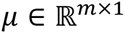 denotes the group-wise expectation of *x_ij_*, and where 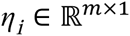 and 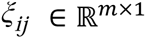 denote within- and between-subject fluctuations, respectively. The random vectors *η_i_* and *ξ_ij_* are assumed to be independent and have expectation zero and covariance matrices Σ*_w_* and Σ*_b_*, respectively (see Section 2.7 for more details). As we will see below, we can express all properties of the similarity indices *R_w_* and *R_b_* and their estimators 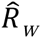 and 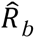 in terms of the model parameters *μ*, Σ*_w_* and Σ*_b_*.

Instead of considering *R_w_* and *R_w_* it will be convenient to consider their Fisher-transformations, denoted by *z_w_* and z*_b_*, respectively, and similarly for their estimators. We first consider the special case in which Σ*_b_* and Σ*_w_* are diagonal matrices (but see panel B of figure **5** to see the observed values of Σ*_b_* and Σ*_w_*), that is 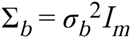 and 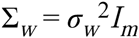 for certain *σ_b_* and *σ_w_* In Section 2.6 it is shown that in this case, the similarity indices can be expressed in terms of the model parameters as follows:

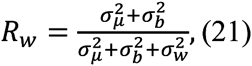

and

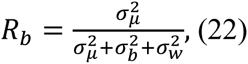

where we have defined 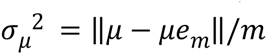. Note that 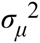 is the variance of FC over links that is common to all subjects. These formulas allow interpreting the similarity indices and relating them to the link-wise ICCs, or more exactly to the parameters determining it.

#### 3.3.2 Quality of functional fingerprints

With respect to the number of sample points (or number of sessions), 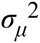 and 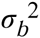 (≈0.05 and ≈0.008 and, respectively) are constants, but 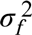 and 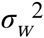 decreases rapidly (the exact speed is irrelevant for now) toward zero, as we showed in figure 4. So, the asymptotic value of *z_b_* = arctanh(*R_b_*) is a limited value (arctanh is the inverse hyperbolic tangent), but as *R_w_* tends to 1, *z_w_* = arctanh(*R_w_*) tends to infinity. Moreover, the variances of *z_b_* and *z_w_* are bounded (by one over the number of links), as the links are correlated between them and the correlations are increasing tending to one. The result is mainly based on the behavior of the sources of variability shown in figure 4, in particular the fact that for every brain *i*, exists a constant FC*_i_*, and that the genuine variability rapidly fade out for increasing number of sample.

To conclude, the model proposed to describe the whole FC is qualitatively in agreement with the experimental results, and it has the advantage of being simple: Few parameters and marginal assumptions determine it completely. In a nutshell: the distributions of (Fisher transformed) *R_w_* and *R_b_* are two Gaussian distributions, whose variances are approximately constant and whose expected values are moving away from one another tending toward infinite values, and with a speed that follows approximately the number of samples.

## 4 Discussion

In this study, we assessed the variability and test-retest reliability of the human functional connectome taking advantage of a unique data set comprising multiple (42) fMRI scans of 5 minutes each for 5 subjects during a classical resting-state paradigm, together with another sets of single-scans obtained from 50 different subjects.

### Single link reliability

From this analysis we obtained the reliability of the functional links between ROIs, as quantified by the ICC. In order to avoid potential biases due to the parcellation, ROIs were obtained both using an anatomical (AAL) as well as a functional parcellation recently proposed by Shen et al. [2013]. From our results we conclude that the average reliability of single link FC is quite low (≈0.2) (figure 3), which is in agreement with the literature (Shehzad et al. [2009]; Birn et al. [2013]). These results, as well as all other results, are qualitatively equal for the two parcellations. Interestingly, we found that the correlation values of all links have an ICC drawn from the same distribution. In other words, our data contains no evidence for heterogeneity of test-retest reliability over links. Indeed, this result suggests an overall homogeneity in the reliability of links in the whole brain, in contrast to what is claimed in the literature (Shehzad et al. [2009]; Zuo and Xing [2014]).

A small ICC variance is crucial to distinguish between reliable and unreliable links. To obtain a small ICC variance, a very large number of the product of subjects and scan sessions is needed (a large number of sessions for few subjects, or vice-versa are equivalent in this sense). To date, analyses of resting-state fMRI test-retest reliability typically used a large number of subjects performing two or three scans; we adopted the opposite strategy, but still did not reach a better power resolution for the ICC. To be more specific, in our data-set we have 42 scans for 5 subjects, and the SD of the estimated ICC was approximately 0.2; similar SD resulted for the data-set analyzed by Zuo and Xing [2014], with 75 subjects and 3 scans. Therefore, to substantially decrease ICC variance in successive studies, we have to use a much higher number of subjects or scans. For example, a fifth of the SD can be achieved with more than 100 participants instead of 6, and 42 scans. These numbers point out the experimental difficulty in differentiating between the reliability of each link of the FC matrix.

### Sources of variability

Thanks to our analysis based on surrogate data, we were able to characterize and quantify different sources of inter-session variability of the correlations between distinct brain regions. As a first approximation, we identify two sources: 1. the statistical uncertainty produced by calculating correlations from a finite number of samples (finite-sample variability), 2. the genuine session-dependent fluctuation of the correlations between different brain regions (within-subjects variability). In order to describe the subject-to-subject variability appropriately, we further identified a third source of variability next to these two sources of variability, that we called simply between-subject variability.

The importance of being able to separate these different sources of FC variability is that it allows us to understand more in depth the temporal dynamics as well as the link-to-link differences in reliability itself. For example, our between-subject variability shows time-consistency, in contrast to the behavior of the finite-sample and within-subject variances, as both decrease for increasing number of sample points. While the decrease of the finite-sample variance with the number of samples is trivial, neither the between- nor the within-subject variances behavior can be foreseen from previous analyses.

Moreover, from this result, and the results of Birn et al. [2013], we can predict that within-subject variance reaches a plateau, at ≈0.012. Indeed, Birn and colleagues showed that the link reliability reaches a maximum value (0.4) for scan duration of approximately 20 minutes. Such a low ICC value limits the use of single link FC as a potential biomarker. Here, we showed that a possible solution can be that of using joining multiple sessions: Our results indicate that in order to obtain an intermediate to high ICC, 6–8 of 5 minutes each sessions are required. Clearly, it would be convenient to use longer scan sessions to diminish the number of scan sessions.

As we showed, the variability of the FC is in part due to the finite sample. Thanks to our surrogate-based analysis, we could quantify how large the relative contributions of the finite sample and genuine variability are. We stress that this genuine (within-subject) variability is much higher than the one reported in Laumann et al. [2015, 2016]. Even though we are not sure what causes this discrepancy, our result seems in quantitative agreement with what has been reported in other reliability studies (Shehzad et al. [2009]; Birn et al. [2013]).

Note that even though the macro-regions do have approximately the same low values of ICC, there is a small region-to-region variability. In principle, these differences can be caused by the three variances; however we showed that the between-subject variance is mainly responsible of the observed structure in ICC. For example, we have described higher values of ICC in cerebellum compared to the lower values of ICC in the pre-frontal region. Similar analysis were carried out in Laumann et al. [2015]; Mueller et al. [2013], however, in those studies the three sources of variability were not completely separated, which makes the results more difficult to interpret.

### From from link-wise unreliability to whole brain stability

We analyzed the similarity between entire FC between subjects and between sessions, describing the consistency of the entire correlation structure within a general linear model (for similar approaches see Mueller et al. [2013]; Finn et al. [2015]). With this statistical model, we provide theoretical ground to understand and solve an apparent paradox: how is it possible that very low link-wise reliability (Birn et al. [2013]) can generate such high stability at the global level (whole FC), as has recently been shown (Finn et al. [2015])?

In particular, we studied the distribution of two similarity indices, *R_w_* and *R_b_*, that measure the distance (in terms of correlation) between FC of two sessions of the same subject and of two subjects, respectively. Taking advantage of the multiple sessions of our data set, we calculated the distribution of these indices for an increasing number of concatenated sessions. In addition, we obtained an approximate expression for the average and the variance of the estimators of the distributions of *R_w_* and *R_b_* (see eqs.18 and 19). These estimators are simple functions of the between- and within-subject variances. In this context, it is straightforward to show that for an increasing number of sessions, the average *R_w_* goes to infinity, while the average *R_b_* remains finite; while their variances are limited. Therefore, if the two distributions do not overlap, the identification is perfect.

Finn et al. [2015] analyzed the identification, without directly adopting these similarity indices. Moreover, they used an increasing number of data-points within the same scan session instead of multiple sessions. The latter is a considerable difference, and as we mentioned before, based on the analysis of Birn and colleagues, we predict a lower asymptotic value for the variance within-subject for scan sessions longer than 20 minutes. This implies, following our analysis, that *R_w_* has an asymptotic limit (upper bound) for scan duration greater than 20 minutes. This prediction is confirmed by the results shown in figure 3B in Finn et al. [2015].

We believe that the relevance of this analysis goes beyond this result: Indeed, we hope that this simple statistical framework can be used as an ordinary tool for further analyzing the FC, and to generate a link between the analysis of the single link and the analysis of the whole FC, or even macro-regions.

### Limitations

The resting-state literature has proposed several measures to characterize spontaneous fMRI fluctuations (see for example the review by Zuo and Xing [2014]). These measures can be related to single voxels Zuo and Xing [2014]), to larger functional networks (based, for example, on independent component analysis), or to the statistical interdependencies between the time-courses of different voxels or regions.

In this study we only focused on one measure, namely the Pearson correlation coefficient obtained from the BOLD signals of different pairs of ROIs. We considered this measure as a starting point, and indeed all analyses performed here can be in principle applied to the measures mentioned above. This choice is motivated essentially by two factors: its simplicity, a linear measure of the relationships between activities, and its widespread use, probably the most commonly used. However, in the recent past different measures of BOLD activities have been presented (see the ones analyzed in Zuo and Xing [2014]) increasing the potentiality of fMRI studies. Here, we choose to study thoroughly the FC generated from the Pearson correlation measure, at the cost of neglecting these other measures.

In our study, we used two parcellations: one anatomical, the AAL, and one based on functional parcellation, proposed in Shen et al. [2013]. In our study, we did not found strong quantitative differences in the results of the two parcellations. However, different studies (e.g., the graph study Fornito et al. [2010]), illustrated the relevance of the parcellation and then we hope to see in future studies an analysis applied to multiple and different kind of parcellations.

In this study, we assessed FC variability without directly analyzing its origin (apart from head-motion). Other studies already started to focus on this important aspect, that can have a very broad application, going from physiological (body heat, cardiac and respiration artifacts, head motion) to technical (machine noise, scanner type, experimental instructions, data standardization, data pre-/post-processing strategies) to brain status (e.g., Birn [2012]; Yan et al. [2013]; Hurlburt et al. [2015]; Yan et al. [2013]; Power et al. [2012]; Laumann et al. [2016]). It would be useful to capitalize the description developed in this work, and to use these insight when planning future studies, therefore to improve our understanding of the sources of variability in the human functional connectome.

The potential of resting-state functional connectivity is well illustrated by its ability to characterize both healthy and abnormal cognitive processes and to predict perception and performance. Further drawing from its potential, however, requires a systematic assessment of its variability and test-retest reliability. Our study has demonstrated how such an assessment, together with the application of appropriate statistical concepts, helps to explain the apparent contradiction between local unreliability and global stability of resting-state fluctuations in the human brain.

## Acknowledgments

This research is supported by the European Research Council (ERC) Advanced Grant DYSTRUCTURE (n. 295129), by the Spanish Research Project PSI2013-42091; PSI2016-75688-P; by the Catalan Agency for Management of University and Research Grants, AGAUR (2014SGR856); (RGB) FI-DGR scholarship of the Catalan Government through the Agencia de Gestío d’Ajuts Universitari i de Recerca, agreement no. 2013FI-B1-00099.

